# A high-throughput platform for feedback-controlled directed evolution

**DOI:** 10.1101/2020.04.01.021022

**Authors:** Erika A. DeBenedictis, Emma J. Chory, Dana Gretton, Brian Wang, Kevin Esvelt

**Author notes:** Designates equal-contribution (Emma J. Chory, Dana Gretton).

## Abstract

Continuous directed evolution rapidly implements cycles of mutagenesis, selection, and replication to accelerate protein engineering. However, individual experiments are typically cumbersome, reagent-intensive, and require manual readjustment, limiting the number of evolutionary trajectories that can be explored. We report the design and validation of Phage-and-Robotics-Assisted Near-Continuous Evolution (PRANCE), an automation platform for the continuous directed evolution of biomolecules that enables real-time activity-dependent reporter and absorbance monitoring of up to 96 parallel evolution experiments. We use this platform to characterize the evolutionary stochasticity of T7 RNA polymerase evolution, conserve precious reagents with miniaturized evolution volumes during evolution of aminoacyl-tRNA synthetases, and perform a massively parallel evolution of diverse candidate quadruplet tRNAs. Finally, we implement a feedback control system that autonomously modifies the selection strength in response to real-time fitness measurements. By addressing many of the limitations of previous methods within a single platform, PRANCE simultaneously enables multiplexed, miniaturized, and feedback-controlled continuous directed evolution.

## INTRODUCTION

Continuous directed evolution enables rapid population diversification and selection by coupling a single desired protein function to fitness, and has been used to engineer biological systems that are not tractable with rational design alone.^1–5^ The most well-established continuous evolution method, Phage-Assisted Continuous Evolution (PACE),^6^ has been successfully applied to a diverse set of protein engineering challenges, including the evolutions of proteases with altered substrate specificity,^7^ of proteins with improved soluble expression,^8^ and of CRISPR-Cas proteins with broadened PAM recognition.^9,10^ The power of this technique comes from the ability to perform many rounds of directed evolution in a single day, making it ideal for challenging evolution goals that require numerous sequential mutations.^11^

Despite its successes, three major challenges exist with current continuous directed evolution methods. First, experiments have a high failure rate. In PACE, phage bearing the evolving biomolecule will frequently “wash out” of the experiment, indicating that selection pressure was too high, or will fail to acquire mutations that improve its activity, indicating that selection pressure was too low. In theory, such systems could substantially benefit from real-time monitoring and feedback control, where selection pressure would be automatically adjusted according to population fitness.^12,13^ Second, experiments are difficult to multiplex, making it challenging to explore enough conditions to observe optimal evolution. In particular, the outcome of an evolution depends upon both the initial genotype^14–16^ and on the experimental conditions of the evolution, including the mutagenesis level,^17^ selection strength,^17,18^ and the order in which different selective pressures are applied.^19^ Third, even when evolution successfully produces an evolved biomolecule, the stochasticity of evolution renders it difficult to derive generalizations or underlying principles from these experiments with limited replicates.^19^ As such, much effort has been spent developing new multiplexed evolution platforms.^20–22^

To comprehensively address these challenges, we developed phage-and-robotics-assisted near-continuous evolution (PRANCE), an automated platform for high-throughput directed evolution capable of activity-dependent feedback control. Whereas traditional phage-assisted continuous evolution (PACE) houses a single evolving population of M13 bacteriophage in a sealed flask,^6^ referred to as a “lagoon,” PRANCE features individual evolving populations contained in the wells of a 96-well plate, enabling highly multiplexed evolution. We validated this platform by performing a well-understood polymerase evolution^6,17,19^ in 96-replicate to characterize its evolutionary stochasticity. We next demonstrated that the miniaturization of lagoons in PRANCE enables the conservation of precious small molecules nearly a hundred-fold by evolving aminoacyl-tRNA synthetases to incorporate non-canonical amino acids. To establish the capacity for multiplexed evolution to evolve a variety of initial genotypes, we ran simultaneous evolutions seeded with variants of 20 different tRNA homologues to evolve quadruplet-decoding tRNAs. Finally, we used an integrated plate reader to implement a feedback control system in which the selection pressure was automatically regulated in response to a real-time luminescence readout of population fitness, which eliminated experiment failure due to phage washout. Overall, PRANCE enables experiments that were once laborious, failure-prone, and severely underpowered to be executed robustly *en masse*.

## RESULTS

### Development of an automated evolution platform

To develop PRANCE, we began by implementing an evolution protocol on a liquid handling robot by placing up to 96 lagoons on-deck that are diluted with fresh bacterial culture twice per hour, enabling each 500 μL experiment to experience 1 vol/hr of media flow-through, comparable to traditional continuous flow PACE (Fig. 1a). The method is initiated by sterilizing and rinsing a 3D-printed 96-well bacterial culture reservoir (Supplementary Fig. 1), which is then filled from either a turbidostat or chemostat in a 37°C incubator, or from a pre-prepared bacterial stock kept in an off-deck 4 °C refrigerator (“cool-PRANCE”) (Fig. 1b). Accessory molecules (e.g., small-molecule inducers, non-canonical amino acids) are then added to the reservoir. We programmed the robot in Python to precisely time the distribution of bacteria, the addition of accessory molecules, the sampling of wells for real-time monitoring, and the disposal of outgoing waste samples (Fig. 1c). Additionally, we built error-handling, failure-mode prevention, and wireless experimenter communication into every PRANCE method (Supplementary Fig. 2) and also optimized the throughput and hands-off time of the robot, such that experimenter intervention is only required once every 7 to 14 hours (Supplementary Fig. 3a,b).

**Fig. 1.**
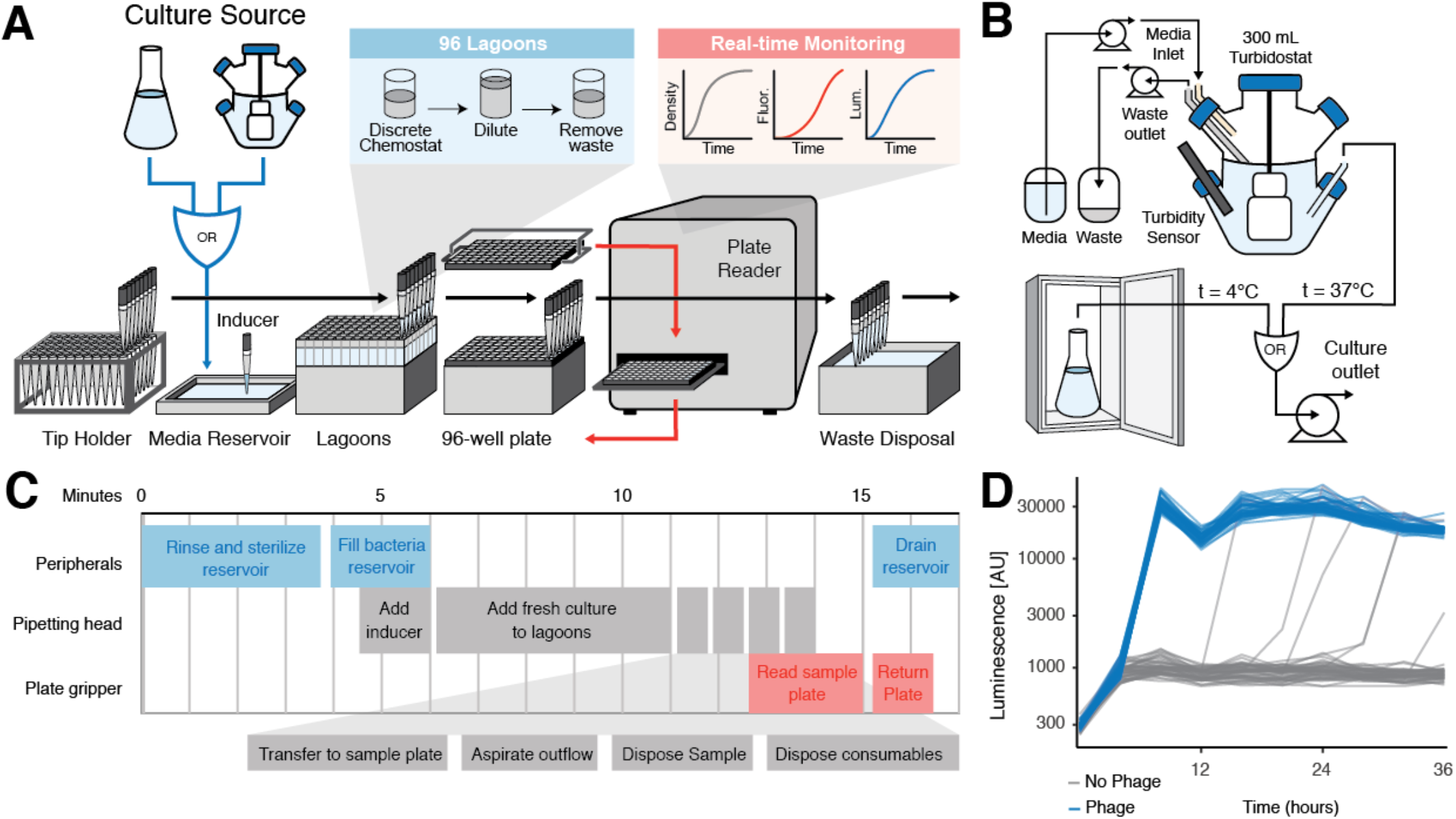
Design and validation of PRANCE. **a,** Real-time monitoring PRANCE method summary for up to 96 simultaneous evolutions. Evolving populations are housed in wells of deep 96-well plates on the deck of a liquid handling robot. Liquid handling is used to create a discrete chemostat for every lagoon, continuously refreshing each population by adding fresh bacterial culture from an off-deck turbidostat and removing waste. An integrated plate reader is used to monitor luminescence or fluorescence readouts of population activity on the evolution goal. Movements by robotic pipette (black arrow) and plate reader (red arrow) movements are shown. **b,** Culture source fluidics (media, turbidostat/static culture, waste) are peripherally separated from the robot. **c,** Asynchronous timing of robotic peripherals in each iteration operate in a loop. **d,** Real-time monitoring of T7 RNAP-expressing phage propagation by monitoring activity-dependent luminescence monitoring of 48 phage lagoons and 48 no-phage lagoons. Each no-phage lagoon that spikes in luminescence represents a cross-contamination event. The cross-contamination rate is measured to be 4 cross-contaminations per 96-lagoons per day.

A key distinguishing feature of PRANCE from other automated evolution platforms^20–22^ is that it enables real-time monitoring of not only bacterial density, but also fluorescence and luminescence within each lagoon. To do so, our method samples the lagoons in discrete intervals, measures these samples in a plate reader, and then either disposes of or preserves the samples for downstream analyses such as plaque assays, sequencing, or other activity measurements of the evolved biomolecule (Fig. 1a). A fluorescent or luminescent reporter protein can be directly coupled to either the presence of phage or to the activity of the evolution circuit itself, enabling real-time monitoring of biomolecule activity. We first demonstrated the real-time monitoring capability of the system using bacteria expressing an M13 phage coat protein (pIII) and a luminescence reporter (luxAB) under the control of a T7 promoter (P_T7_). We then inoculated 96 total lagoons both with or without T7 RNA polymerase (T7 RNAP)-expressing bacteriophage and demonstrate successful phage-detection in all 48 phage-induced samples in less than 4 hours (Fig. 1e). In traditional PACE experiments, phage plaque assays are commonly used to monitor titer over the course of the experiment. However, plaque assays are laborious and require overnight incubation which delays the ability to course-correct the evolution conditions of any given lagoon. While in-line luminescence monitoring has previously been implemented with PACE,^23^ burdensome equipment requirements have limited its adoption. In this work, we demonstrate that luminescence data strongly correlates with activity-dependent plaque assay-determined phage titers, allowing us to track evolution progress in real-time (Supplementary Fig. 4a). Although less correlative, absorbance depression—the decrease in a lagoon’s OD_600_ resulting from slowed bacterial growth due to phage infection—can also be used to track phage propagation (Supplementary Fig. 4b), providing a real-time monitoring strategy for any phage-based evolution in which a reporter protein cannot be readily inserted within the selection. Overall, we found real-time monitoring data to be invaluable for interpreting the results of highly multiplexed evolutions.

Phage contamination is also known to be problematic when attempting parallel evolutions with PACE; thus, we sought to develop a reliable method of measuring cross-contamination between samples and identify a robotic configuration that minimizes it. We found that using a 96-channel head exhibits minimal cross-contamination (Fig. 1e), compared to an 8-channel pipette (Supplementary Fig. 3c) as measured by luminescence observed in no-phage control wells, at a rate of 4 cross-contaminations per 96-lagoons per day. Thus, this approach also allows us to, for the first time, quantify the incidence of cross-contamination problems in phage-assisted evolutions. An additional important factor in PRANCE success is host bacteria because phage infection occurs via the F pilus, which *E. coli* express during mid-log phase.^24^ As with traditional PACE, PRANCE can source host bacteria from a chemostat (bacteria cultured at constant flowthrough rate) or turbidostat (bacteria cultured at constant turbidity). However, we also discovered that PRANCE can utilize bacteria grown to mid-log phase and then refrigerated, eliminating the requirement for continuous bacterial culture entirely (Supplementary Fig. 3d). The ability to incorporate chilled cultures in PRANCE has broad downstream utility, as it enables easy use of numerous types of bacterial cultures and may ultimately be the most user-friendly mode of PRANCE operation.

### Multiplexing enables investigation of evolutionary dynamics

While current directed evolution methods may suffice when the primary goal is to obtain a functional protein, their application to the investigation of evolutionary dynamics has been limited by the irreproducibility of directed evolution experiments and the difficulty of running many experiments in parallel. As proof of principle, we sought to characterize the fitness landscape for the previously-described evolution of the T7 RNAP to initiate using a new promoter.^6,17,19^ We prepared host bacteria expressing pIII and luxAB under the control of the T3 promoter (P_T3_) on an accessory plasmid (AP) and containing a mutagenesis plasmid (MP) promoting the greater sampling of genetic diversity during evolution^25^ (Fig. 2a). Next, we inoculated 96 total lagoons with or without T7 RNAP-expressing M13 phage lacking pIII (90 experimental lagoons with 6 no-phage controls) and tracked their evolution in real time with absorbance depression (Fig. 2b) and luminescence (Fig. 2c). This high-throughput exploration of the evolution of the T7 RNAP allowed us to measure the frequencies with which different genotypes arose across replicate evolution experiments, with both novel and previously described mutations having been observed (Fig. 2e). In addition, we examined the distribution of elapsed times until the acquisition of those genotypes, which we found to be approximately normally distributed (Fig. 2d). Thus, high-replicate evolutions enabled by PRANCE allow for the thorough characterization of an evolutionary fitness landscape in a single experiment, a task that would have been infeasible with traditional directed evolution techniques.

**Fig. 2.**
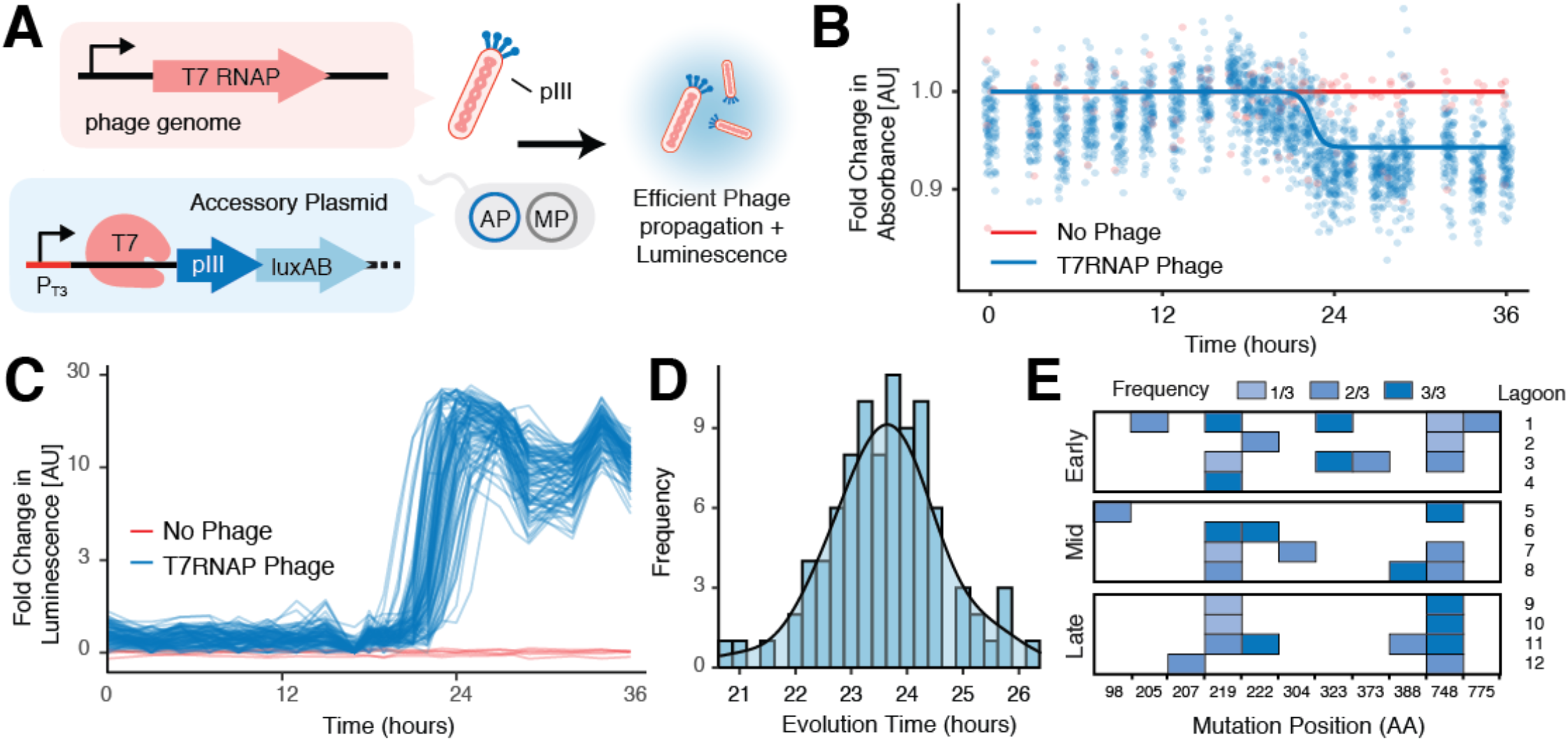
High-throughput directed evolution. **a,** Strategy for evolving T7 RNAP to recognize the T3 promoter, in which expression of the required pIII protein is driven by the T3 promoter on an accessory plasmid and the evolving T7 RNAP is encoded on the replicating phage genome. **b,** Real-time absorbance depression monitoring of 90 simultaneous evolutions with 6 no-phage controls, fit with a binomial regression of the total data. **c,** Real-time luminescence monitoring of 90 simultaneous evolutions with 6 no-phage controls. **d,** Time required to acquire mutations permitting the T7 RNAP to recognize the T3 promoter, obtained from binomial regressions of the total data (Supplementary Fig. 5). Smoothed fit is calculated with a kernel density estimate. **e,** Mutations observed from 12 representative lagoons at early (−1σ), mid (mean), late (+1σ) time points. Mutational frequency refers to n number of phage that were sequenced, 3 per lagoon.

### Miniaturization enables reagent-limiting evolutions and the routine inclusion of controls

Despite the many successful applications of PACE to diverse classes of proteins, its widespread adoption has been limited by the large quantities of reagents and media that it requires for extended periods of evolution—a feature that is especially prohibitive for applications which require expensive, novel, or difficult-to-synthesize small molecules. In addition, the difficulty of multiplexing with PACE discourages the routine inclusion of control lagoons that would assist the interpretation of experimental results. PRANCE reduces the total reaction volume of each lagoon by 100-fold, making small-molecule-intensive applications feasible while enabling the extensive use of controls. To demonstrate these capabilities, we modified an established evolution of aminoacyl-tRNA synthetases (AARSs) for the installation of non-canonical amino acids (ncAAs)^26^ with PRANCE while using minimal amounts of ncAAs.

First, we encoded both a pyrrolysyl-tRNA synthetase (PylRS) and a UAG-tRNA^Pyl^ within the M13 phage genome, while encoding an amber codon-containing pIII and luciferase reporter on an AP in the host cells, similar to previously described^26^, with the modification that encoding the tRNA on the phage enables greater multiplexing (Fig. 3a). Thus, phage proliferation and luminescence are directly coupled to amber suppression of the UAG stop codon upon incorporation of an ncAA (Fig. 3b). We validated that three synthetase variants—a chimeric PylRS (chPylRS), an evolved variant exhibiting intermediate activity for incorporation of the ncAA Boc-lysine (chPylRS-IP), and a highly-active variant identified upon extended evolution times (chPylRS-IPYE)—demonstrated increasing kinetic activities of Boc-lysine incorporation by measuring luminescence upon induction of AARS expression with IPTG (Fig. 3c). Additionally, decreased phage propagation is observed when additional UAG stop codons are incorporated into pIII, as expected (Fig. 3d).

**Fig. 3.**
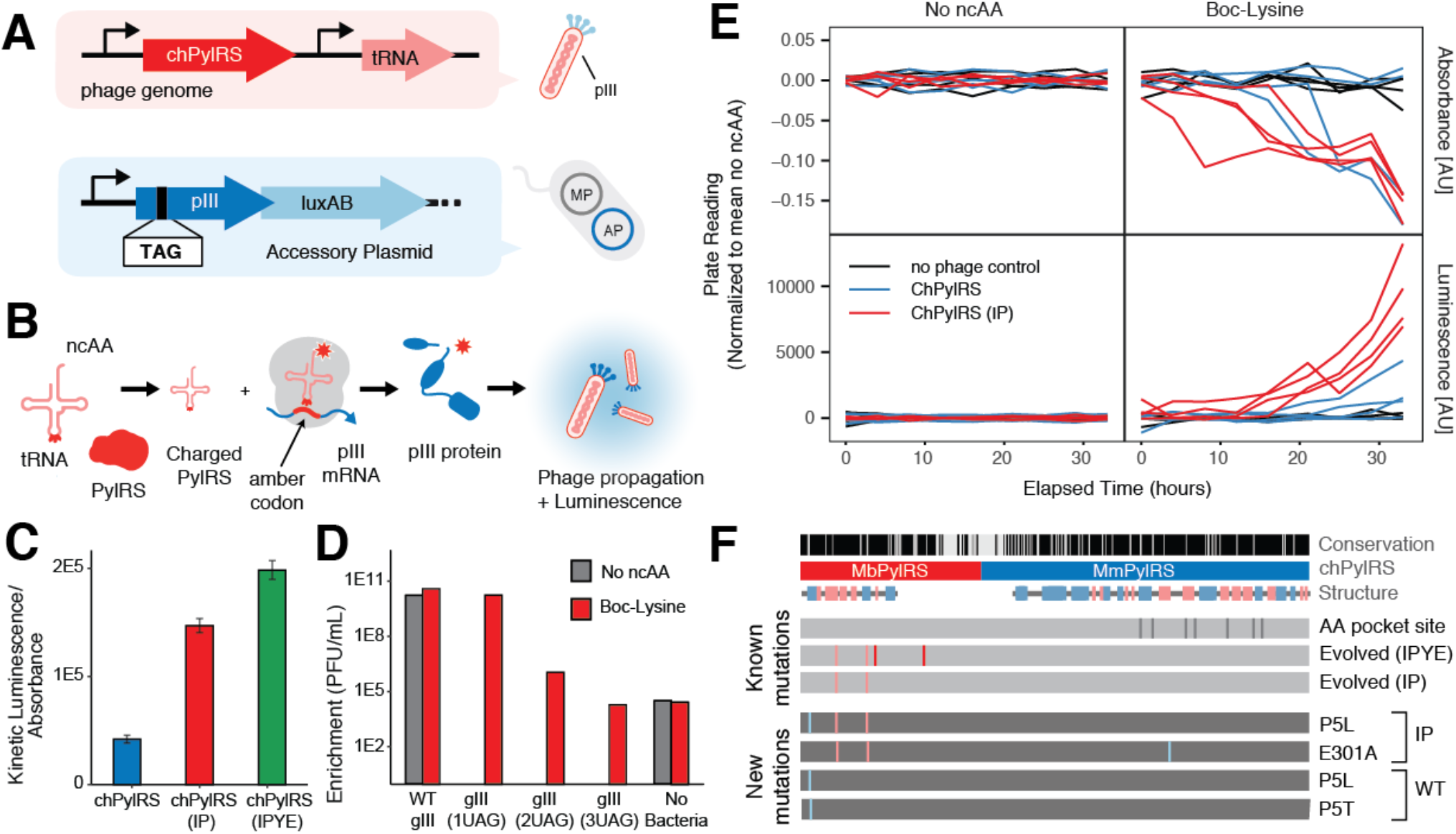
Multiplexed directed evolution of AARSs. **a,** Strategy for evolving AARSs and tRNAs to incorporate non-canonical amino acids. TAG amber codons are inserted into the pIII protein, and the evolving tRNA^pyl^ and chPylRS are encoded in the evolving phage genome. **b,** Phage propagation and luminescence are contingent on the successful incorporation of ncAAs into pIII. **c,** Efficiency of unevolved versus evolved chPylRS variants (IP, IPYE mutations) at incorporating Boc-lysine into a luxAB reporter, normalized to no-ncAA and no-IPTG controls. **d,** Quantifying the selection stringency of incorporating 1, 2, or 3 ncAAs into pIII, in either the presence or absence of Boc-lysine, using the evolved variant chPylRS-IPYE. **e,** Real-time absorbance depression and luminescence monitoring of chPylRS evolutions, beginning from either the unevolved state (chPylRS, blue) or an intermediate evolved state (chPylRS-IP, red). **f,** Genotypes of the evolved variants. Phage persisted in all of the chPylRS-IP lagoons, and clonal phage acquired novel mutations (E301A or P5L). Only half of the chPylRS lagoons resulted in persistent phage propagation, but each acquired different mutations at the same N-terminal proline residue (P5L or P5T), within the conserved essential N-terminal domain of PylRS.^30,31^

We then used PRANCE to recapitulate the evolutions of chPylRS and chPylRS-IP for Boc-lysine incorporation. We seeded 8 lagoons containing Boc-lysine with phage encoding either chPylRS or chPylRS-IP in quadruplicate, hypothesizing that we would observe the emergence of chPylRS-IPYE upon evolution. In order to characterize whether evolutions of chPylRS or chPylRS-IP were prone to the emergence of “cheater” AARSs that incorporate canonical amino acids,^27^ we also seeded 8 lagoons with phage encoding each chPylRS variant in the absence of ncAAs. Together with 8 no-phage lagoons, we monitored luminescence and absorbance across a total of 24 lagoons over 36 h of evolution time. We observed luminescence of both chPylRS- and chPylRS-IP-encoding phage in the presence of Boc-lysine (Fig. 3e); further analysis revealed that the genotypes of evolved variants included previously unidentified mutations (Fig. 3f). The absence of luminescence in evolution control lagoons lacking Boc-lysine indicate that cheater AARSs are unlikely to emerge under the employed evolution conditions (Fig. 3e). Thus, the inclusion of control lagoons enabled the extraction of additional information previously unobtainable from a single PACE experiment. In addition, this PRANCE experiment involving the continuous addition of Boc-lysine to 12 lagoons consumed less than 100 mg of Boc-lysine, compared to the nearly 1 g that would be required for a single lagoon in a 36 hour PACE experiment. Given this substantial reduction in reagent costs, PRANCE may enable the multiplexed and well-controlled evolutions of AARSs to install ncAAs that are either too expensive (e.g., 4-azido-Phe, $2500/g^28^), rare (e.g., pyrrolysine^29^), or novel to be easily used in traditional PACE.

### Multiplexing enables the simultaneous evolution of dozens of quadruplet-decoding tRNAs

Previously, we demonstrated that PACE can be used to evolve quadruplet-decoding tRNAs (qtRNAs)^32,33^ as part of a larger effort to engineer a translation system entirely based on four-base codons that could incorporate multiple noncanonical amino acids in a single protein^34^ (in revision). However, the success of qtRNA evolution can differ drastically depending on which tRNA homologue is used to seed the evolution, and on which of the possible quadruplet codons is targeted. Thus, performing sufficient experiments to identify a set mutually compatible qtRNAs for all 20 canonical amino acids as well as numerous non-canonical amino acids would be exceedingly challenging with traditional PACE. Here, we used PRANCE to identify functional AGGG qtRNAs by subjecting a full set of 20 different tRNA homologues to evolution in a single experiment.

We began by replicating the evolution of six TAGA-decoding qtRNAs with PRANCE, and observed similar results and evolved genotypes as evolution performed with traditional PACE equipment (in revision) (Supplementary Fig. 6). Next, we inserted an AGGG quadruplet codon into pIII and encoded a variety of qtRNAs into the M13 phage genome (Fig. 4a), tying successful quadruplet-decoding to phage replication (Fig. 4b). In the absence of a functional qtRNA, the quadruplet codon generates a frameshift, thereby truncating pIII and precluding phage propagation.

**Fig. 4.**
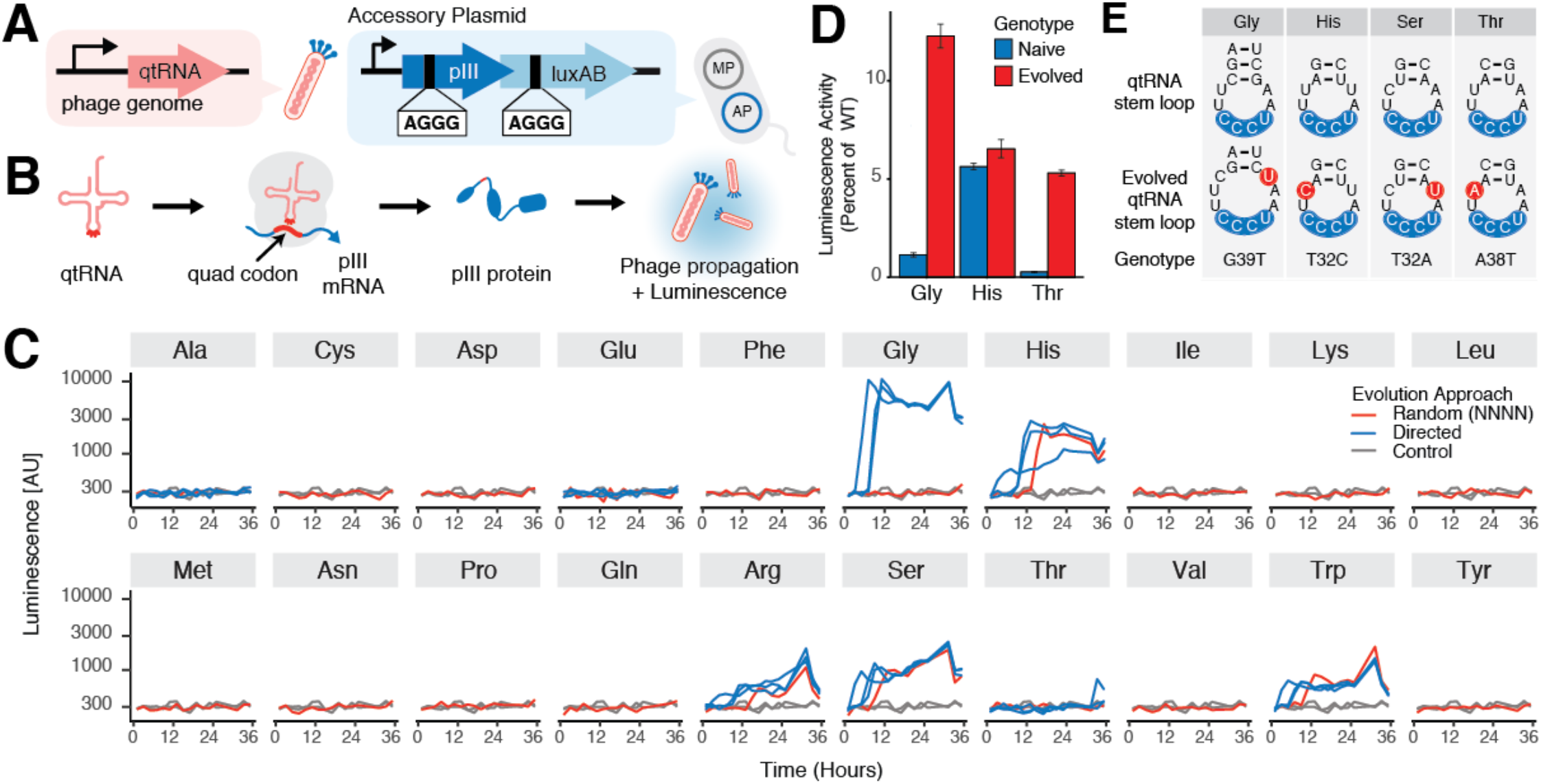
Multiplexed directed evolution of qtRNAs. **a,** Strategy for evolving qtRNA to recognize quadruplet codons. AGGG! quadruplet codons are inserted into the pIII protein and luxAB, and the evolving qtRNA are encoded in the evolving phage genome. **b,** Phage propagation and luminescence are contingent on the successful decoding of the quadruplet codons in pIII and luxAB proteins respectively. Failure to decode a quadruplet codon results in premature termination and truncated protein. **c,** Real-time absorbance and luminescence monitoring of qtRNA-encoding phage where either randomized or directed anticodons were used. **d,** Efficiency of evolved versus unevolved qtRNA at incorporating amino acids compared to a triplet codon. Evolved Ser-AGGG T32A is too toxic to be subcloned. **e,** Initial and evolved qtRNA genotypes.

Finally, we initiated PRANCE by seeding 1) 20 lagoons with phage encoding 20 different qtRNA homologues corresponding to every canonical amino acid, each containing a library of randomized anticodons (NNNN); 2) 24 lagoons with phage encoding 8 different qtRNA homologues (for Ala, Glu, Gly, His, Arg, Ser, Thr, and Trp), each with directed !AGGG anticodons, in triplicate; and 3) 4 lagoons consisting of evolution controls. Altogether, we observed evolutionary outcomes across 48 lagoons in a single experiment.

Luminescence traces indicated that phage encoding Gly, His, Ser or Thr qtRNA homologues successfully evolved (Fig. 4c); indeed, when subcloned, several of the evolved qtRNAs exhibited improved activity (Fig. 4d). In addition, while we largely detected no difference in evolutionary outcome between qtRNAs with randomized or directed anticodons, we observed that Gly-qtRNAs with a directed anticodon rapidly evolved, in contrast to Gly-qtRNAs with randomized anticodons, which did not support phage propagation. This result confirms the importance of seeding evolutions with diverse genotypes in order to maximize the probability of achieving engineering goals. Thus, with PRANCE we were able to evolve multiple AGGG-decoding qtRNAs in a single experiment, an effort that would have previously involved dozens of individual PACE experiments.

### Feedback control enables real-time modulation of stringency

Though we successfully identified evolved AGGG-decoding qtRNAs with PRANCE, we also observed a high failure rate. Many of the qtRNAs with low initial activity (Ala, Cys, Asp, etc.) never evolved, and instead the phage washed out of the lagoons. Additionally, qtRNAs with especially high initial activity (Arg, Trp) persisted in the lagoons and triggered luminescence monitoring, but did not acquire any mutations (Fig. 4c). These results indicate that selection was too stringent for some, and too lenient for others. We hypothesized that applying feedback control to dynamically tune the selection stringency in response to the measured activity of each lagoon could greatly improve the likelihood of successfully evolving biomolecules from diverse, challenging starting points.

In order to test the efficacy of feedback control on continuous directed evolution, we implemented a feedback control system that adjusts the stringency of selection in response to a real-time luminescence readout of biomolecule activity. We hypothesized that this capability would allow us to 1) evolve multiple biomolecules side-by-side that differ greatly in initial activity, 2) reduce the possibility of phage washout, and thereby evolution failure, and 3) maintain selection pressure even after active variants of the initial genotypes have been found in order to discover further improved mutants. As a model system, we selected four qtRNA homologues (Phe, His, Ser, and Arg) that decode TAGA. All four are known to have evolved variants, but differ greatly in initial activity (Supplementary Fig. 6). Previously, obtaining the highest-activity variants of all four qtRNAs required sequential rounds of continuous flow with different bacterial APs (in revision). Thus, we aimed to implement feedback control to enable the extended evolution of all four qtRNAs in a single experiment.

First, we characterized three bacterial APs that confer different levels of selection pressure (Fig. 5a). The “easiest” of these APs encodes T7 RNAP with two quadruplet codons incorporated, with the T7 promoter driving production of pIII and luxAB. Circuit architectures containing T7 RNAP are frequently used when the initial activity of the biomolecule is very low because production of a single functional polymerase has an amplifying effect.^7,8,35^ We also characterized more stringent APs containing one (“medium”) or two (“hard”) quadruplet codons in pIII. We found that these three APs provide good coverage of the phage enrichment space that corresponds to the initial activity of these qtRNAs (Fig. 5a).

**Fig. 5.**
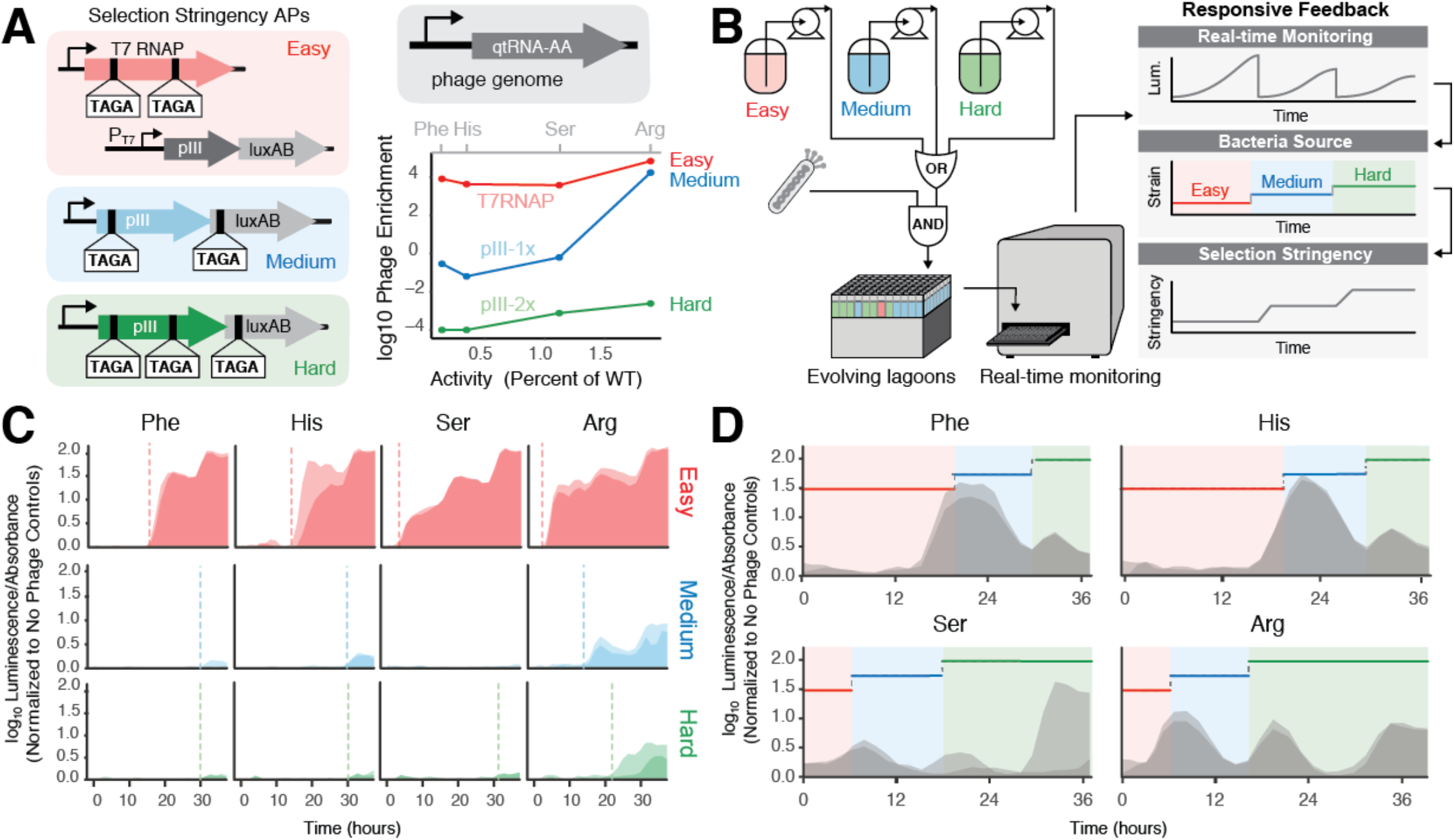
Feedback control of continuous evolution. **a,** Strategy for evolving qtRNAs with increasing stringency of selection (red/easy = T7 RNAP, medium/blue = pIII-1x, hard/green = pIII-2x). The transfer function of each AP’s stringency enables evolution of qtRNAs at different starting activities (Phe, His, Ser, Arg). Starting activity is quantified as “Percent of WT”, or the luminescence generated by co-expressing the qtRNA and luxAB-357-TAGA, relative to an all-triplet luxAB. **b,** Three bacterial strains with increasing AP stringencies are added to evolving lagoons and monitored in real time. The bacteria source for each well is adjusted in response to real-time luminescence measurements with feedback control to automatically increase stringency of evolving phage populations. **c,** Evolutions of 4 qtRNAs on APs with varying stringencies. Dotted lines indicate first emergence of phage. **d,** Real-time luminescence measurements of evolving qtRNAs with feedback-controlled selection stringency intervals.

First, we propagated each of the four qtRNA-encoding phages on each bacterial source in duplicate. We found that all four can stably propagate on the “easy” bacteria to varying degrees; however, only the Arg-qtRNA, which has the highest initial activity, can support phage propagation on the more stringent bacterial sources (Fig. 5c). Next, we allowed the robotic system to control which of these three bacteria types are designated as the source culture for each individual lagoon (Fig. 5b). To do so, we implemented a simple feedback adjustment strategy in which the bacterial source is changed in response to an increase in normalized luminescence (see Methods). This strategy successfully propagated phage encoding all four types of qtRNAs through the end of the 36-hour experiment, at which point all lagoons were being serviced with the most stringent bacterial source (Fig. 5d). Thus, feedback-controlled PRANCE avoided phage washout in all cases while simultaneously exposing high-activity qtRNAs to more challenging evolution conditions. These results demonstrate that PRANCE comprises a robust solution to the high failure rate of continuous directed evolution experiments.

## DISCUSSION

We report the development of PRANCE, a high-throughput platform for continuous directed evolution. PRANCE is designed to accommodate multiplexed evolution, minimize reagent and media volumes, enable periodic sampling of the lagoons for downstream analysis, monitor evolution progress in real time, and implement feedback control for within-experiment modulation of selection stringency on a lagoon-by-lagoon basis. We adapted PRANCE to source bacteria from either a continuous culture stirred turbidostat/chemostat or cool standing culture, enabling evolution experiments with a variety of experimental requirements. Multiplexed evolution with PRANCE addresses two key weaknesses of directed evolution experiments in that it enables evolutions without knowing which initial genotypes and experimental conditions will yield the most productive result *a priori*, and provides a solution for minimizing or quantifying experiment-to-experiment variability. We have demonstrated how PRANCE can be used to 1) evolve T7 RNAP to recognize the T3 promoter in 96-replicate, 2) evolve AARS/tRNA pairs requiring only small quantities of accessory small molecules, 3) evolve 20 different qtRNA homologues simultaneously, and 4) implement feedback control to evolve multiple qtRNAs with diverse initial activities while avoiding phage washout. Altogether, these efforts encompassing >100 evolving populations required only four independent PRANCE experiments.

Though the current method is presently consumable-intensive when operating at maximum capacity (96 lagoons), further method development could greatly extend the hands-off user window above our standard operating time frames of 7, 14 or 21 hours by implementing tip cleaning cycles. While currently designed primarily for phage-assisted continuous evolution, it should also be feasible to extend the system to continuous diversification platforms such as those used with yeast^2,22^ and mammalian cells^3,4^ when utilized within a sterilized, laminar flow environment. In the future, PRANCE could be implemented to perform previously infeasible evolution experiments such as the characterization of complex fitness landscapes, the evolution of small molecule-dependent proteins, and experiments integrating responsive negative selections—experiments facilitated by the PRANCE-specific features of multiplexing, volume miniaturization, and feedback control, respectively.

PRANCE can uniquely perform feedback control triggered by real-time measurements of biomolecule activity during evolution. In our examples, we utilized activity-dependent luciferase measurements quantified by an integrated plate reader. However, any measurement taken using robot-integrated hardware could potentially be used to trigger feedback control in PRANCE, such as qPCR, ELISAs, or even orthogonal cell-based screens. Additionally, we have shown that a simple feedback controller can be used to greatly reduce experiment failure—a key advance of immediate practical utility to the field. In the future, the feedback-control capability will enable experimental testing of control strategies for molecular evolution^12^ to optimize evolutionary outcomes.

Directed evolution has long been a discipline in which experimental outcomes have been determined by chance. PRANCE enables researchers to readily traverse a substantial fraction of the possible fitness landscape within a single, well-controlled, and responsive experiment. Our hope is that by eliminating the time-consuming guesswork of such experiments, PRANCE can help to reinvent directed evolution as a more reliable and reproducible engineering discipline.

## Acknowledgements

We would like to acknowledge Alvaro Cuevas and Hamilton Robotics for their guidance and assistance in implementing a python API. We thank Kristala Prather’s laboratory for equipment use and assistance. We thank Ethan Alley and Stephen Von Stetina for their thoughtful comments on the manuscript.

## Author contributions

E.A.D. and K.M.E. conceived the study. E.A.D. and D.G. developed the platform with advice from K.M.E. D.G. developed the software with advice from E.A.D. and K.M.E. E.A.D., E.J.C., and D.G. designed the experiments with advice from K.M.E. E.A.D, E.J.C., D.G, and B.W. performed the experiments. E.J.C., E.A.D, and B.W. wrote the manuscript with input from all authors.

## Supplementary Figures

**Supplementary Fig. 1.**
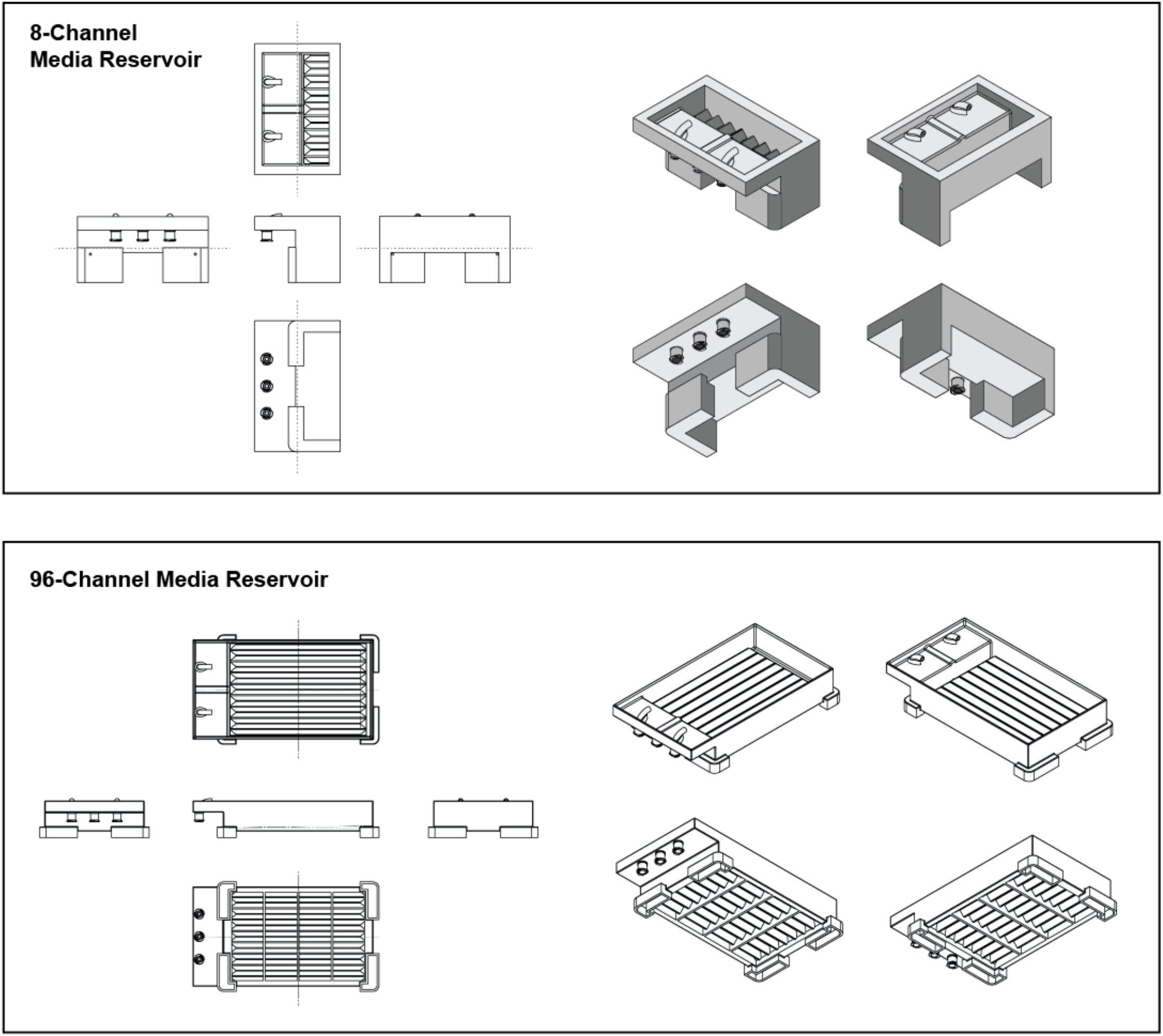
Reservoir diagrams. Schematics of the 8-channel and 96-channel media reservoirs. These were printed on a Form 3 resin 3D printer. See the Extended Supplement for .stl files for each.

**Supplementary Fig. 2.**
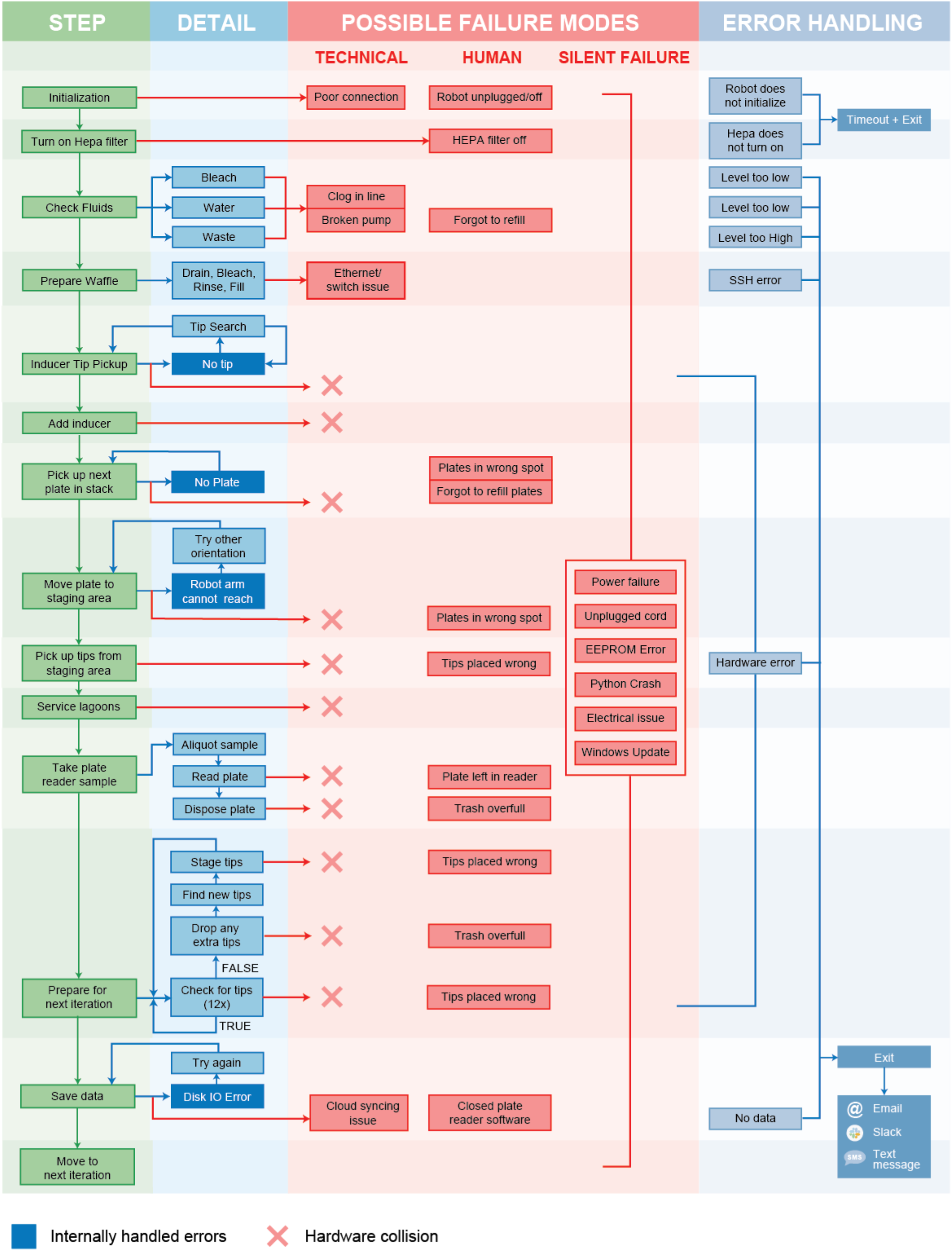
Failure mode analysis.

**Supplementary Fig. 3.**
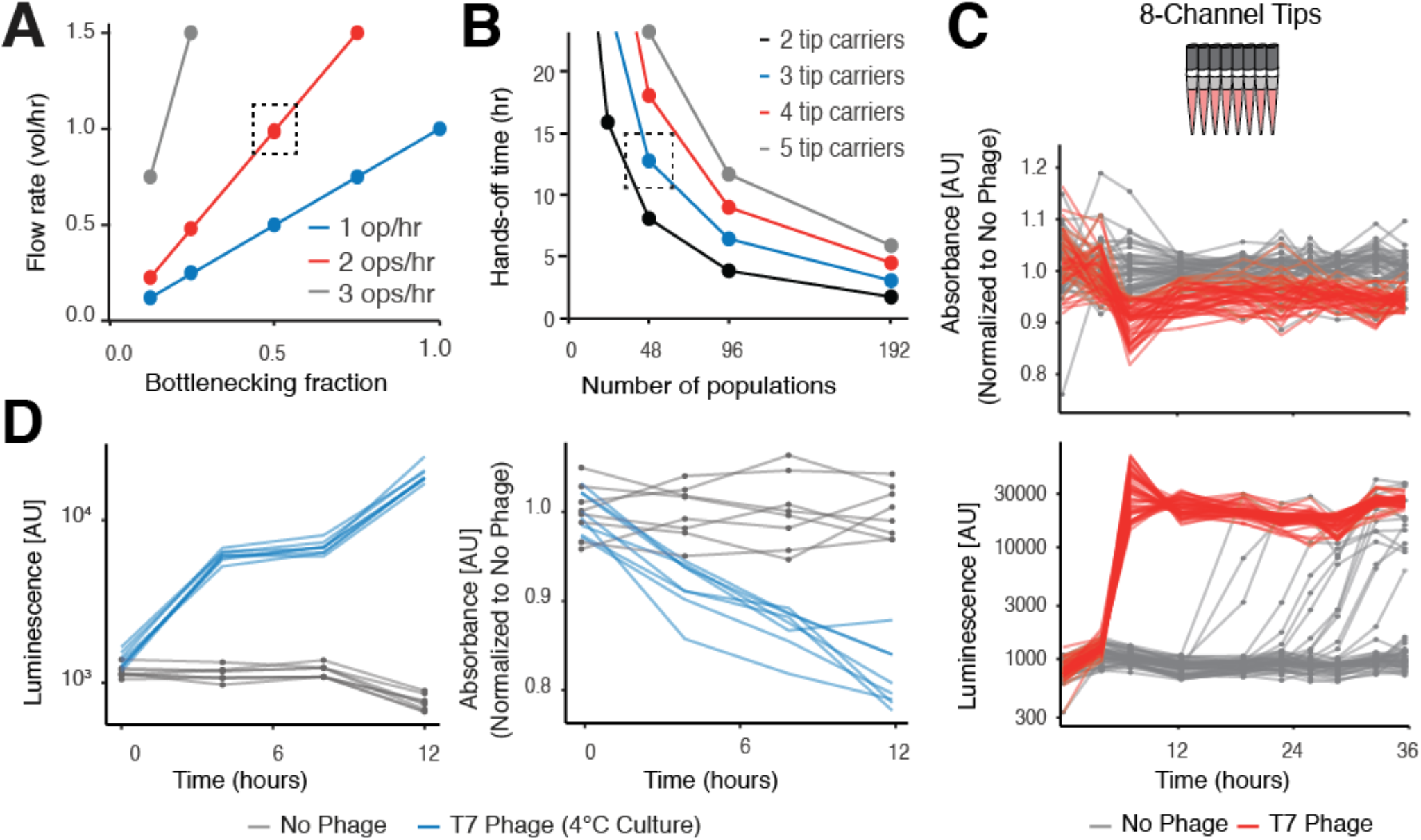
PRANCE optimization. **a,** The maximum flow-through rate is determined by the frequency with which the robot exchanges liquid (operations per hour), as well as the fraction of the standing volume of the population that is exchanged during each operation (the bottlenecking fraction). There is a trade-off between the maximum flow rate and the extent of bottlenecking. **b,** The number of populations that can be serviced assuming 2 ops/hr impacts the hands-off time between adding more consumables to deck. Larger robot decks can fit more tip carriers, more tips, and therefore require less frequent servicing. **c,** Real-time monitoring of T7 RNAP-expressing phage propagation by monitoring absorbance depression and luminescence of 48 phage lagoons and 48 no-phage lagoons serviced with the 8-channel pipetting head introduces more cross-contamination than the 96-channel head due to increased flyover events (see Fig 1d). **d,** Real-time monitoring of T7 RNAP-expressing phage propagation from a bacteria source stored at 4 °C by monitoring absorbance depression and luminescence.

**Supplementary Fig. 4.**
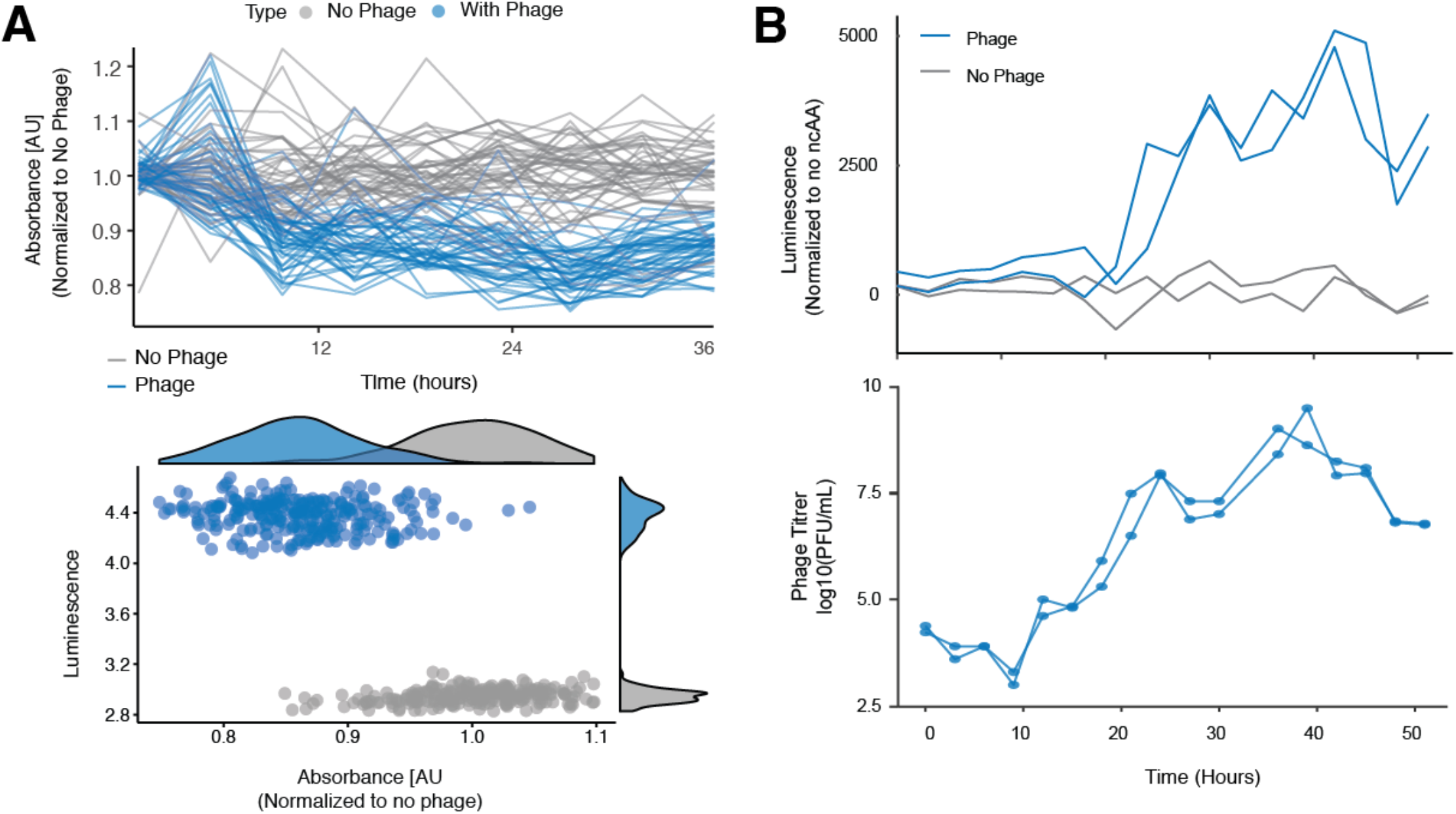
Relationship between real-time monitoring data and phage titer. **a,** Correlation between absorbance depression and luminescence. Luminescence data from Figure 1d. **b,** Comparison of luminescence and phage titer as measured by traditional plaque assays.

**Supplementary Fig. 5.**
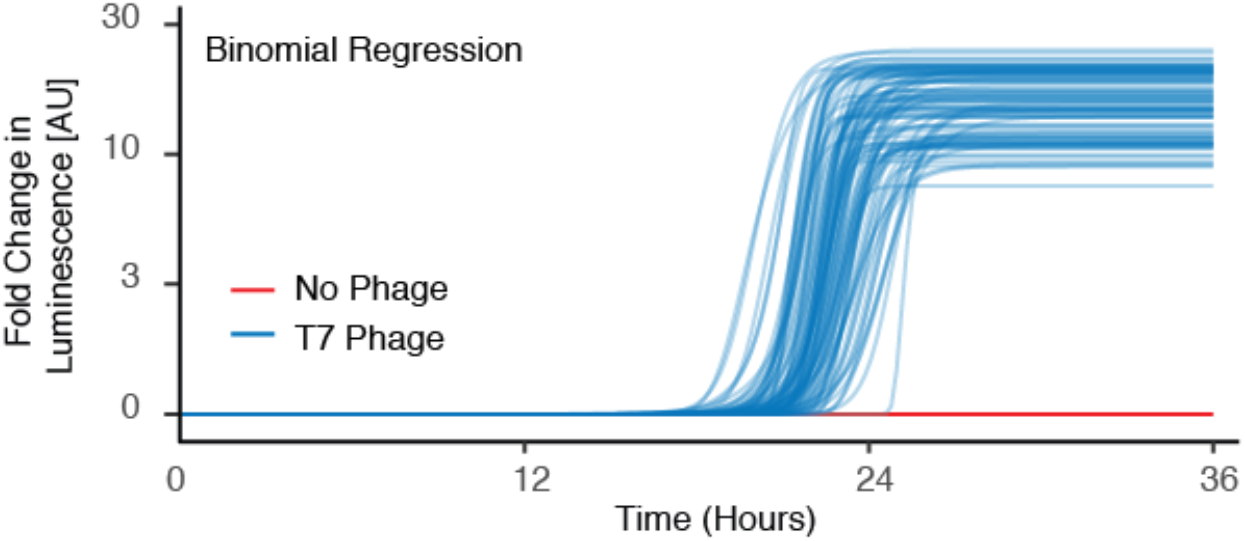
Binominal regression of luminescence data during T7 RNAP evolution to bind the T3 promoter.

**Supplementary Fig. 6.**
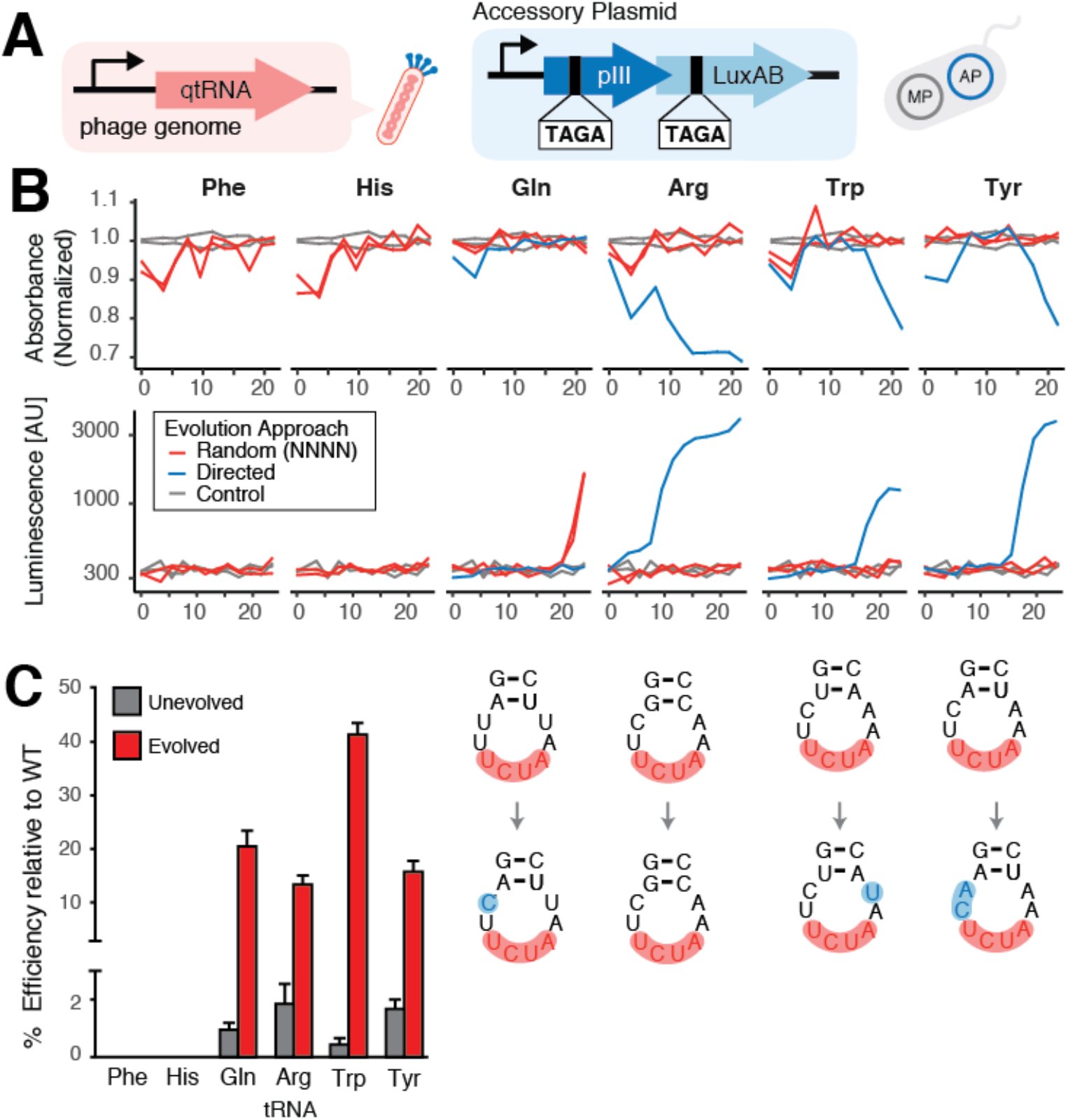
TAGA-qtRNA PRANCE. **a,** Constructs for evolving TAGA-decoding qtRNAs. **b,** Representative results for evolving TAGA-decoding qtRNAs. **c,** Evolved qtRNAs exhibit increased ability to decode a TAGA quadruplet codon. Units are the percent luminescence when translating luxAB-357-TAGA in the presence of the qtRNA relative to expression of all-triplet-luxAB.

**Supplementary Fig. 7.**
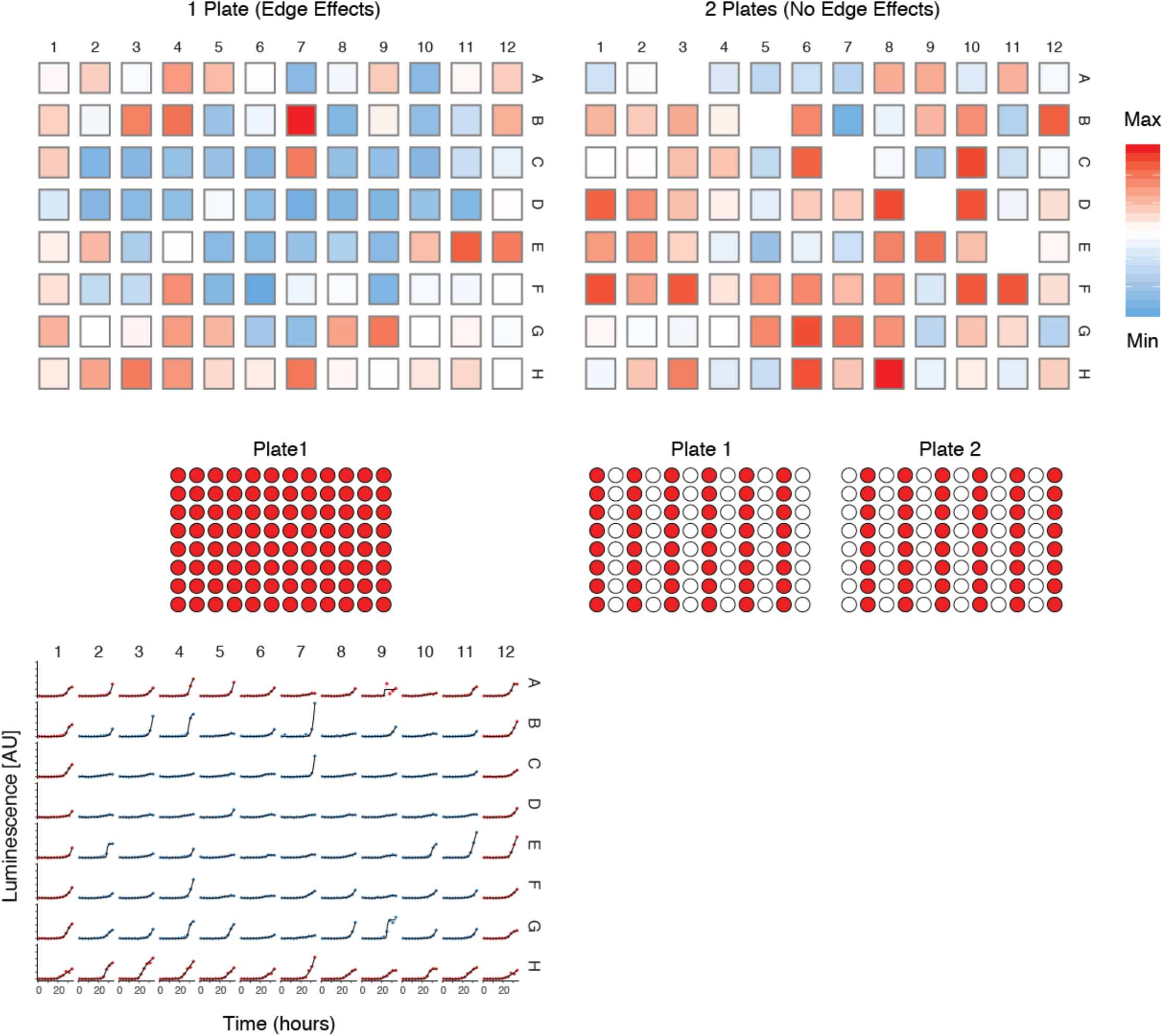
Edge effects. Comparison between 96 lagoons implemented in a densely packed 96-well plate (left) and 96 lagoons split over two plates to reduce edge effects (right). Data plots the minimum time to phage detection via luminescence monitoring (below). Both plates above are normalized to their internal max value.

**Supplementary Table 1.** Method details for each experiment. In all cases, the flow-through rate was 1 vol/hr. MP6^25^ was used as the mutagenesis plasmid in all evolution experiments. See the Extended Supplement [<u>link</u>] for plasmid maps.

| Experiment | Figure | Bacterial source | Lagoon layout | AP/Phage |
| --- | --- | --- | --- | --- |
| Phage propagation with 96-head | SI Fig 7 (left) | Turbidostat | 96 lagoons in a dense 96-well plate. All wells inoculated with phage. | AP: pJC173b. Phage: WT T7 RNAP selection phage * |
| Phage propagation with 96-head | SI Fig 7 (right) | Turbidostat | 96 lagoons, every-other-column layout, spread over two plates. All wells inoculated with phage. | AP: pJC173b. Phage: WT T7 RNAP selection phage |
| Phage propagation with 96-head | Figure 1d, SI Fig 4a, SI Fig 7 (right) | Turbidostat | 96 lagoons, every-other-column layout, spread over two plates. Checkerboard phage inoculation pattern. | AP: pJC173b. Phage: WT T7 RNAP selection phage |
| Phage propagation with 8-channel head | SI Fig 3c | Turbidostat | 96 lagoons, every-other-column layout, spread over two plates. Checkerboard phage inoculation pattern. | AP: pJC173b. Phage: WT T7 RNAP selection phage |
| Phage propagation with chilled bacteria | SI Fig 3d | Chilled bacteria | 16 lagoons, every-other-column layout. | AP: pJC173b. Phage: WT T7 RNAP selection phage |
| T7 → T3 RNAP evolution | Figure 2, SI Fig 5 | Turbidostat | 96 lagoons, every-other-column layout, spread over two plates. | AP: pJC174b*. Phage: WT T7 RNAP selection phage |
| AARS evolution | Figure 3, SI Fig 4b | Turbidostat | 24 lagoons, every-other-column layout. | AP: pEC182a<br>Phage: ECSP1x11, ECSP1x13 ** |
| AGGG qtRNA evolution | Figure 4 | Turbidostat | 48 lagoons, every-other-column layout. | AP: pED15x7, Phage: all 20 ESP1x phage libraries and !AGGG phage variants |
| TAGA qtRNA evolution | SI Fig 6 | Turbidostat | 48 lagoons, every-other-column layout. | AP: pED15x1, Phage: all 20 ESP1x phage, and phage libraries |
| Feedback control qtRNA evolution | Figure 5 | Chemostats | 48 lagoons, every-other-column layout. | Easy AP: pED7x163 + pED20x5. Medium AP: pED15x1 + ES rope. Hard AP: pED15x2 + ES rope. Phage: ESP1xR-1, ESP1xS-1, ESP1xH-1, ESP1xF-1. |

\* pJC173b, pJC174b, and WT T7 RNAP selection phage were gifts from Ahmed Badran.
\*\* chPylRS AP and phage derived from pDB038a and pTECH-chPylRS. pTECH-chPylRS(IPYE) was a gift from Dieter Söll (Addgene plasmid # 99222 ; http://n2t.net/addgene:99222 ; RRID:Addgene\_99222). pDB038a was a gift from David Liu (Addgene plasmid # 99211 ; http://n2t.net/addgene:99211 ; RRID:Addgene\_99211).

## METHODS

See the Extended Supplement [<u>link</u>] for part numbers, and all plasmids used in these experiments.

### General methods

#### Antibiotics

Antibiotics (Gold Biotechnology) were used at the following working concentrations: carbenicillin, 50 μg/mL; spectinomycin, 100 μg/mL; chloramphenicol, 40 μg/mL; kanamycin, 30 μg/mL; tetracycline, 10 μg/mL; streptomycin, 50 μg/mL.

#### Media

Davis Rich Media (DRM)^26^ is used for PACE and all experiments involving plate reader measurements due to its low fluorescence and luminescence background. 2XYT media, a media optimized for phage growth, is used for all other purposes, including phage-based selection assays and general cloning. See Extended Supplement for media catalog numbers.

#### Preparation of chemically competent cells

Strain S2060, a K12 derivative optimized for directed evolution^36^ was used in all luciferase, phage propagation, and plaque assays, as well as in all PACE experiments. To prepare competent cells, an overnight culture was diluted 1,000-fold into 50 mL of 2XYT media supplemented with maintenance antibiotics and grown at 37 °C with shaking at 230 rpm to OD_600_ ~0.4–0.6. Cells were pelleted by centrifugation at 6,000x *g* for 10 min at 4 °C. The cell pellet was then resuspended by gentle stirring in 5 mL of TSS (LB media supplemented with 5% v/v DMSO, 10% w/v PEG 3350, and 20 mM MgCl2). The cell suspension was stirred to mix completely, aliquoted and flash-frozen in liquid nitrogen, and stored at −80 °C until use. See Extended Supplement for catalog numbers.

#### Transformation of chemically competent cells

To transform cells, 100 μL of competent cells were thawed on ice. To this, plasmid (2 μL each of miniprep-quality plasmid; up to two plasmids per transformation) and 100 μL KCM solution (100 mM KCl, 30 mM CaCl2, and 50 mM MgCl2 in H2O) were added and stirred gently with a pipette tip. The mixture was incubated on ice for 10 min and heat shocked at 42 °C for 90 s. The mixture was chilled on ice for 4 min, then 850 μL of 2XYT media was added. Cells were allowed to recover at 37 °C with shaking at 230 rpm for 0.75 h, streaked on 2XYT media + 1.5% agar plates containing the appropriate antibiotics, and incubated at 37 °C for 16–18 h. See Extended Supplement for catalog numbers.

#### Phage supernatant filtration

To filter 500 μL of phage, bacteria were pelleted by centrifugation at 8,000x *g* for 2 min in a tabletop centrifuge. Supernatant was transferred to a 0.22 μm filter column, and centrifuged at 1000x *g* for 1 min to create filtered phage flowthrough. To filter 50 mL of phage supernatant, 50 mL of culture was similarly pelleted. Supernatant was applied to a Steriflip 0.22 μm vacuum filter unit. To filter up to 150 μL of phage in 96-well plate format, the 96-well plate of bacteria was pelleted by centrifugation at 1,000x *g* for 10 min. 150 μL of supernatant was applied to wells of a 96-well 0.22 μm filter plate taped atop a 96-well PCR plate, and centrifuged at 1000x *g* for 1 min to create filtered phage flow-through. Phage can be stored at 4 °C in 96-well plate format covered with an aluminum sealing film. For frequently-accessed phage samples, we recommend storage in 2 mL screw cap tubes in order to minimize potential phage contamination generated from opening snap-caps. See Extended Supplement for catalog numbers.

#### Standard phage cloning

Competent *E. coli* S2060 cells were prepared (as described) containing pJC175e, a plasmid expressing pIII under control of the phage shock promoter.^11^ To clone ΔpIII M13 bacteriophage, PCR fragments were assembled using USER as above. The annealed fragments were transformed into competent S2060-pJC175e competent cells (as described), which complement pIII deletion from the bacteriophage. Transformants were recovered in 2XYT media overnight, shaking at 230 rpm at 37 °C. The phage supernatant from the resulting culture was filtered (as described), and plaqued (as described). Clonal plaques were expanded overnight, filtered, and Sanger sequenced. See Extended Supplement for catalog numbers.

#### tRNA diagrams

R2R was used to generate tRNA diagrams. R2R is free software available from http://www.bioinf.uni-leipzig.de/~zasha/R2R/.

### Plaque assays

#### Manual Plaque assays

S2060 cells were transformed with the Accessory Plasmid (AP) of interest. Overnight cultures of single colonies grown in 2XYT media supplemented with maintenance antibiotics were diluted 1,000-fold into fresh 2XYT media with maintenance antibiotics and grown at 37 °C with shaking at 230 rpm to OD_600_ ~0.6–0.8 before use. Bacteriophage were serially diluted 100-fold (4 dilutions total) in H2O. 100 μL of cells were added to 100 μL of each phage dilution, and to this 0.85 mL of liquid (70 °C) top agar (2XYT media + 0.6% agar) supplemented with 2% Bluo-Gal was added and mixed by pipetting up and down once. This mixture was then immediately pipetted onto one quadrant of a quartered Petri dish already containing 2 mL of solidified bottom agar (2XYT media + 1.5% agar, no antibiotics). After solidification of the top agar, plates were incubated at 37 °C for 16–18 h. See Extended Supplement for catalog numbers.

#### Robotics-accelerated plaque assays

The same procedure was followed as above, except that plating of the plaque assays was done by a liquid handling robot (Hamilton Robotics) by plating 20 μL of bacterial culture and 100 μL of phage dilution with 200 μL of soft agar onto a well of a 24-well plate already containing 235 μL of hard agar per well. To prevent premature cooling of soft agar, the soft agar was placed on the deck in a 70 °C heat block. Source code from our implementation can be found at https://github.com/dgretton/roboplaque

### Phage enrichment assays

S2060 cells were transformed with the Accessory Plasmids (AP) of interest as described above. Overnight cultures of single colonies grown in 2XYT media supplemented with maintenance antibiotics were diluted 1,000-fold into DRM media with maintenance antibiotics and grown at 37 °C with shaking at 230 rpm to OD_600_ ~0.4–0.6. Cells were then infected with bacteriophage at a starting titer of 10^5^ pfu/mL. Cells were incubated for another 16–18 h at 37 °C with shaking at 230 rpm. Supernatant was filtered (as described) and stored at 4 °C. The phage titer of these samples was measured in an activity-independent manner using a plaque assay containing *E. coli* bearing pJC175e (as described). A complete list of catalog numbers can be found in the Extended Supplement.

### Equipment set-up for PRANCE

#### General robotic equipment and configuration

A Hamilton Microlab STARlet 8-channel base model was augmented with a Hamilton CORE 96 Probe Head, a Hamilton iSWAP Robotic Transport Arm, and a Dual Chamber Wash Station. Air filtration was provided by an overhead HEPA filter fan module integrated into the robot enclosure. A BMG CLARIOstar luminescence multi-mode microplate reader was positioned inside the enclosure, within reach of the transport arm.

#### Mini peristaltic pump array

Up to seven miniature 12 volt, 60 mL/min peristaltic pumps (“fish tank pumps”) were actuated by custom motor drivers. A Raspberry Pi mini single-board computer received instructions over local IP and commanded the motor drivers via I^2^C. A complete list of catalog numbers can be found in the Extended Supplement.

#### Bacterial reservoir

We implemented a self-cleaning bacterial reservoir consisting of the pump array and a custom 3D-printed corrugated reservoir (Supplementary Fig. 1); see Extended Supplement for .stl file. Following each filling of the bacterial reservoir with fresh bacteria and liquid exchange (see below), remaining culture was drained from the reservoir and the reservoir was automatically rinsed with 5% bleach once and water four times. The reservoir is seated on a Big Bear Automation Microplate Orbital Shaker, which shakes the reservoir throughout all self-cleaning steps to achieve uniform diffusion.

#### Software

A general-purpose driver method was created using MicroLab STAR VENUS ONE software and compiled to Hamilton Scripting Language (hsl) format. Instantiation of this method and management of its local network connection was handled in Python. The Pyhamilton Python package provided an overlying control layer. Interfaces to the CLARIOstar plate reader, pump array, and orbital shaker were encapsulated in supporting Python packages. We used Git to develop and version control the packages and the specific Python methods used for each experiment; our software implementation can be found on github at https://github.com/dgretton/std-96-pace

### PRANCE experimental set-up

#### Mutagenesis functionality plating test

In order to avoid premature induction of the mutagenesis plasmid (MP) (which can mutagenize the MP itself and lead to loss of inducible mutagenesis) all bacteria containing MP are repressed with 20 mM glucose, and were not grown to higher density than OD_600_ = 1.0. In order to confirm MP functionality, a bacterial culture sample or four single colony transformants were picked and resuspended in DRM, and were serially diluted and plated on 2XYT media + 1.5% agar with appropriate antibiotics (at half of the usual concentrations), and either 25 mM arabinose or 20 mM glucose. Bacteria containing functional MP6 will exhibit smaller colonies on the arabinose plate relative to the glucose plate.

#### PRANCE host bacterial strain preparation

To prepare host bacterial strains, the Accessory Plasmid of interest and mutagenesis plasmid (MP6)^25^ were transformed into S2060 cells as described. During MP6 transformation, cells were recovered with DRM to ensure MP6 repression, and were streaked on 2XYT media + 1.5% agar with appropriate antibiotics (at half of the usual concentrations), and 20 mM glucose. Transformants were picked for the mutagenesis functionality plating test (see above), and those same transformants were grown at 37 °C with shaking at 230 rpm to OD_600_ 0.4. This culture was used to seed a PRANCE culture in a manner dependent on the specific PRANCE experiment:

- To initiate a *turbidostat*, prepared host bacteria were used to seed DRM supplemented with 25 μg/mL carbenicillin and 20 μg/mL chloramphenicol in a turbidostat (implemented by a standard turbidity probe and glassware vessel). The bacteria were grown with stirring at 37 °C to OD_600_ = 0.8.
- To initiate *chemostats*, prepared host bacteria were used to seed 150 mL of DRM supplemented with appropriate antibiotics (at half of the usual concentrations) in three separate chemostat vessels. The chemostat cultures were kept at ~100–150 mL and grown with stirring at 37 °C. The flow-through and volume of the turbidostats were manually adjusted as needed to maintain the target OD_600_ = 0.8 in all chemostats.
- To prepare *chilled bacterial culture* for use in PRANCE, an overnight culture of prepared host bacteria was diluted 1:1000 into 2 x 600 mL of DRM in two separate 2L baffled flasks with appropriate antibiotics (at half of the usual concentrations). The bacteria were grown at 37 °C with shaking at 230 rpm to OD_600_ = ~0.3–0.4. The two 600 mL cultures were combined and stored at 4 °C for up to 4 days before being situated in a 4 °C refrigerator adjacent to the robot.

MP6 functionality is also confirmed at the end of the experiment with the arabinose/glucose plating test (see above).

#### General PRANCE implementation

Host bacteria were pumped from their source (turbidostat, chemostat, or refrigerated chilled culture) into the on-deck bacterial reservoir. Arabinose was added to the reservoir to a final concentration of 10mM to induce mutagenesis. 250 μL of bacteria were then robotically transferred to each lagoon (500 μL/lagoon) in deep 96-well plates situated on the deck of the liquid handling robot, which was kept at 37 °C. In order to avoid edge effects, lagoons are placed in every other column (Supplementary Fig. 7). This liquid exchange occurs every 30 min continuously throughout the duration of the experiment, generating a flow-through rate of 1 vol/hr in each lagoon. After allowing mutagenesis induction for 1 hr, each lagoon was inoculated with selection phage to a starting titer between 10^5^-10^8^ pfu/mL, with appropriate lagoons set aside as no-phage controls. Sampling and plate reading of lagoon waste liquid was automatically carried out every 1–4 h as described. After completion of PRANCE, the phage supernatant from select lagoons was filtered and plaqued; clonal plaques were expanded overnight, filtered, and Sanger sequenced.

#### Semi-continuous flow using pipetting

To perform liquid exchange, every 30 minutes automated pipetting was used to exchange liquid in on-deck lagoons. The procedure consisted of tip pickup, aspiration of fresh bacterial culture, dispense of new culture into lagoon vessels, mixing of liquid, aspiration of waste, dispense of waste into the waste reservoir, and tip disposal.

#### Automated plate reading

At specified times, 175 μL of the waste samples are deposited into plate reader plates. To read the plates, the plate gripper transports the plate to the integrated plate reader for absorbance at OD = 600 nm and fluorescence or luminescence measurements as required.

#### Choosing number of lagoons

Consult Supplementary Fig. 3 to select a configuration (number of lagoons, bottlenecking fraction, and hands-off time) that meets your experimental requirements. There are two primary limiting factors: the amount of time the robot requires to perform each necessary step within a given time frame, and the number of consumable reagents (tips, plates, etc.) that can be physically staged on the deck of the robot. For every individual PRANCE run, we select and tune the method configurations depending upon the requirements of each individual experiment (Supplementary Table 1). All of our experiments are run at an effective flow-through rate of 1 vol/hr, with a bottlenecking fraction of 0.5. A configuration we frequently preferred consisted of 48 evolutions, requiring experimenter intervention once every 13 hours. Alternatively, when operating at maximum capacity (1 vol/hr, 96 lagoons) PRANCE requires intervention once every 7 hours. The trade-off between lagoons and hands-off time will depend upon the capacity of your robot deck.

#### Feedback controller calculation

For each individual lagoon, luminescence is first normalized to absorbance. Each lagoon reading is then smoothed over an interval of n=3 and stringency adjustment is triggered after 3 sequential events in which the reading is 3σ over background. Background is quantified as the average of no phage controls (luminescence/absorbance). The second stringency adjustment (medium → hard) is triggered similarly, following a 12 hour delay to allow for luminescence equilibration.

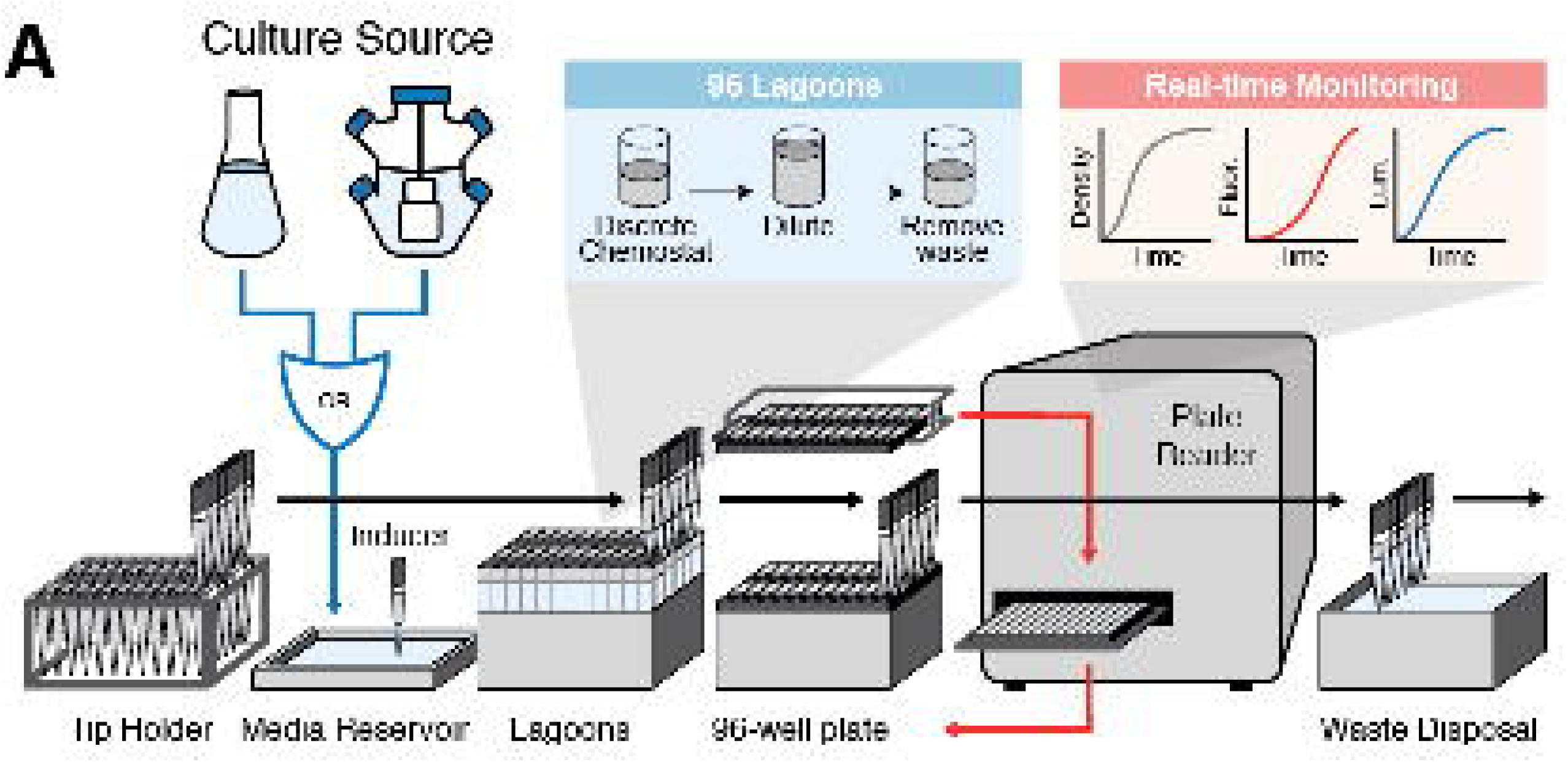

